# IsoSolve: an integrative framework to improve isotopic coverage and consolidate isotopic measurements by MS and/or NMR

**DOI:** 10.1101/2021.03.08.430771

**Authors:** Pierre Millard, Sergueï Sokol, Michael Kohlstedt, Christoph Wittmann, Fabien Létisse, Guy Lippens, Jean-Charles Portais

## Abstract

Stable-isotope labeling experiments are widely used to investigate the topology and functioning of metabolic networks. Label incorporation into metabolites can be quantified using a broad range of mass spectrometry (MS)and nuclear magnetic resonance (NMR)spectroscopy methods, but in general, no single approach can completely cover isotopic space, even for small metabolites. The number of quantifiable isotopic species could be increased, and the coverage of isotopic space improved, by integrating measurements obtained by different methods; however, this approach has remained largely unexplored because no framework able to deal with partial, heterogeneous isotopic measurements has yet been developed. Here, we present a generic computational framework based on symbolic calculus that can integrate any isotopic dataset by connecting measurements to the chemical structure of the molecules. As a test case, we apply this framework to isotopic analyses of amino acids, which are ubiquitous to life, central to many biological questions, and can be analyzed by a broad range of MS and NMR methods. We demonstrate how this integrative framework helps to i) clarify and improve the coverage of isotopic space, ii) evaluate the complementarity and redundancy of different techniques, iii) consolidate isotopic datasets, iv) design experiments, and v) guide future analytical developments. This framework, which can be applied to any labeled element, isotopic tracer, metabolite, and analytical platform, has been implemented in IsoSolve (available at https://github.com/MetaSysLISBP/IsoSolve and https://pypi.org/project/IsoSolve), an open source software that can be readily integrated into data analysis pipelines.

---

Stable-isotope labeling experiments are widely used to investigate metabolic networks in the fields of systems biology^1–2^, biotechnology^3–4^ and biomedical research^5–6^. The most effective approach is to combine ^13^C-labeling strategies with a detailed analysis of isotope incorporation into metabolites, as measured by mass spectrometry (MS) and/or nuclear magnetic resonance (NMR) spectroscopy^7.^ MS provides global isotopic information by quantifying the proportions of molecules with different numbers of tracer isotopes (isotopologue distributions)^8–9^, while NMR provides positional information on tracer incorporation at specific positions in the molecules (isotopomer distributions)^10–13^ by exploiting the ^1^H and ^13^C nuclei via non-decoupled experiments – such as homonuclear ^1^H-^1^H-TOCSY and heteronuclear ^1^H-^13^C-HSQC experiments.

Each separate NMR and MS method provides partial isotopic information by quantifying specific (sets of) isotopic species. MS(/MS) is used to quantify isotopologue distributions of complete molecules or fragments^8–9^, ^14^ where each carbon isotopologue contains several isotopic species that cannot be distinguished since they all have the same mass9. NMR is similarly limited in that it only quantifies a subset of isotopic species since positional information is in general limited to a small part of the carbon skeleton (typically from 1 to 3 carbon atoms depending on the experiment). Recently, ^15^Nand pure-shift-NMR experiments have successfully been applied to access long-range heteronuclear coupling constants, thereby increasing the number of isotopic species that can be quantified^11^, ^13^. Nevertheless, none of the available methods provides complete coverage of isotopic space, even for small metabolites.

Integrating measurements from different approaches should expand the range of quantifiable isotopic species^7^, thus improving the coverage of isotopic space. This is exploited in ^13^C-fluxomics studies, where different datasets are frequently integrated using isotopic models of metabolic networks to improve flux quantification^4^, ^15–16^. However, a major drawback of model-based integrative approaches is their strong dependence on the assumptions and simplifications of the model (e.g. the topology of the metabolic network as defined in the model) and the fact that labeling has to be quantified in several metabolites. As an alternative approach, intuitive reasoning has proven useful in improving the coverage of isotopic space by defining relationships between isotopic measurements directly at the level of the molecule. This has been demonstrated for the combination of two NMR experiments, ZQF-TOCSY and HSQC, which allowed absolute quantification of 4 of the 8 isotopomers of a block of three carbon atoms^10^. A few relationships have also been established between specific MS and NMR datasets^17–18^, but the heterogeneity of the isotopic information obtained by MS and NMR makes integrating measurements from these two platforms difficult. Overall, the lack of a generic integrative framework has meant that the potential expansion of isotopic coverage that could be achieved by combining independent MS(/MS) and/or NMR datasets has remained largely untapped.

In this article, we present a computational framework that can be used to integrate any type of isotopic data. We demonstrate how this framework allows isotopic coverage to be clarified and improved, thereby consolidating isotopic measurements. As a test case, we apply this framework to isotopic analysis of amino acids, which are ubiquitous to life, abundant, and central to many biological questions. They can be analyzed by a broad range of MS and NMR methods so the dataset considered here is representative of the wide range of measurements these analytical platforms can provide.

## EXPERIMENTAL SECTION

### ^13^C-labeled standard of amino acids

The reference material we used to evaluate the proposed methodology was a biologically produced sample containing ^13^C-labeled amino acids with a controlled and predictable isotopic composition: the isotopic species of each amino acid are forced to all be present at the same concentration. The nature, production, and qualification of this standard sample have been described in detail previously^9^, ^19^.

### Isotopic analyses

We analyzed the ^13^C-labeled reference material by GC-MS^20^, and by HSQC^15^ and ZQF-TOCSY^15^ NMR experiments, as detailed in the corresponding publications. We also gathered additional (HNCA, HACO-DIPSY, LC-MS) datasets for this reference material from the literature^11, 19^.

### Implementation of the computational framework

The frame-work developed in this work was implemented as a Python 3 module named IsoSolve. This can be used both as a command-line tool and as a module imported into Python scripts. The intuitive data input is based on tab separated value (tsv) files. The first (mandatory) input file describes the relationships between the measurements and isotopomers as presented in the Results section. The formalism is universal and can be used for all existing and future types of measurement. The second (optional) input file contains the numerical values of the measurements and their associated standard deviations. If provided, these numerical data are used to consolidate the measurements by solving a non-linear least square problem. The symbolic formulas obtained can be verified by assigning randomly drawn values to isotopomers (and thus to the corresponding measurements) and comparing the randomly drawn and calculated values. Explicit results and details of the calculation process can be consulted in a user-friendly HTML document and/or as python variables for later programming use. IsoSolve also generates isotopically enriched InChIs for all isotopic species (https://github.com/MSI-Metabolomics-Standards-Initiative/inchi-isotopologue-extension), facilitating its integration into standardized data analysis pipelines. IsoSolve is freely available under open source license (GPL v2) at https://pypi.org/project/IsoSolve. A Jupyter notebook (https://jupyter.org) is also provided at https://github.com/MetaSys-LISBP/IsoSolve_notebook as an introduction to programming applications of the software. This notebook contains the code used to perform all the calculations in this study and generate all the equations and Figures 3–6.

## RESULTS AND DISCUSSION

### General principle

The essence of the proposed framework lies in the way fundamental relationships between measurements and the underlying isotopic species are exploited. As a rule, isotopic measurements from any method can be expressed as the relative abundance of a (set of) isotopic species in a larger set of species. These relationships can be expressed as a system of equations linking independent measurements through isotopic space. We formalize this principle and illustrate how it can be exploited to integrate any type of measurement using the example of alanine, which is routinely analyzed by ^13^C-NMR, ^1^H-NMR and MS methods.

^13^C-NMR experiments provide information on positional isotopomers through J_CC_ coupling patterns, i.e. on the isotopic content of the carbons bonded to the (labeled and detected) carbon. The J_Cα-CO_ and J_Cα-Cβ_ coupling constants of alanine are typically resolved, so the ^13^C-NMR signal of the C_α_ atom has four components (*a*, *b*, *c*, *d*), which correspond to 4 individual isotopic species (010, 110, 011 and 111, where 0 and 1 refer to ^12^C and ^13^C, respectively, and where the first digit corresponds to the CO group, the second to C_α_ and the third to C_β_). Their abundance is measured relative to the total amount of isotopic species that contribute to these signals, i.e. all species with a labeled C_α_ atom. C_α_ ^13^C-NMR signals can thus be expressed as:

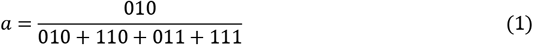

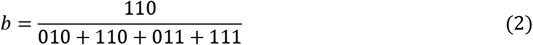

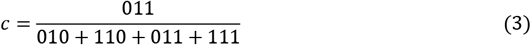

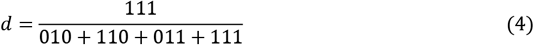

It should be stressed that this definition ensures that the measurements always add up to 1, a fact that is subsequently used to simplify formulas, as detailed below. In practice, this means that after measuring integrated intensities in arbitrary units, the measurements have to be normalized to their sum. This can always be done provided at least one signal is quantified.

^1^H-NMR experiments provide information on specific enrichments via JCH coupling patterns, i.e. on the proportion of ^12^C (*e*) and ^13^C (*f*) isotopes in the carbon bonded to the analyzed proton. The signal of the H_α_ proton can thus be expressed as:

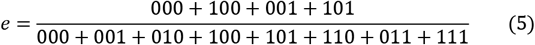

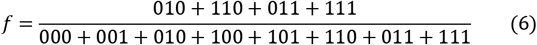

Finally, the total abundance of the set of isotopic species is set by convention to unity, yielding an additional equation:

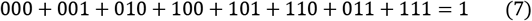

Measurements obtained by ^1^Hand ^13^C-NMR can be integrated by solving this system of equations (eqs 1–7). Expressing the abundance of all isotopic species as a function of the measurements yields the following solution:

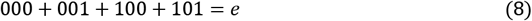

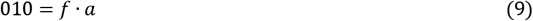

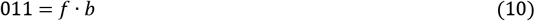

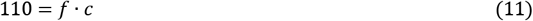

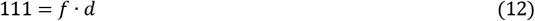

This system is undetermined, in that while the summed abundance of the four species (000+001+100+101) can be calculated from the measurements, their individual abundances cannot. Nevertheless, the integration of ^1^H- and ^13^C-NMR data yields the absolute abundance of four isotopic species (010, 011, 110 and 111), i.e. their abundance is expressed relative to the total amount of molecule rather than to a subset of species (010+110+011+111). This information is new from an analytical standpoint because it cannot be obtained from individual experiments but only by integrating them, as described previously^10^.

In contrast to NMR, MS distinguishes molecular entities in terms of the number of labeled atoms incorporated, i.e. by distinguishing between isotopologues. This information can be obtained for different elementary metabolite units (EMUs), which are defined as moieties comprising distinct subsets of the compound's atoms^17^. The carbon isotopologue distribution (CID) is the vector of isotopologue abundances of a given EMU, where the abundance of each isotopologue is expressed relative to the total amount of molecule. The CID of the EMU containing all the carbon atoms of alanine is represented by a vector [*g*, *h*, *i*, *j*] and is formally defined by the following equations:

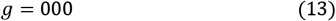

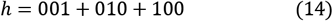

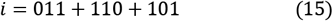

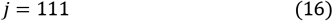

MS thus provides four additional equations (eqs 13–16) which can be combined with those derived from the ^1^Hand ^13^C-NMR data (eqs 1–7). Solving this extended system of equations yields:

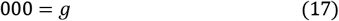

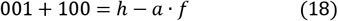

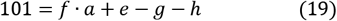

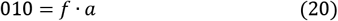

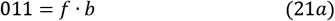

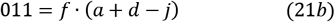

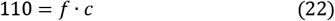

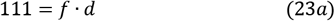

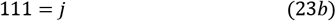

Integrating MS data provides new information, namely the absolute abundances of species 000, 101, and the summed abundance of 001 plus 100. The two latter species cannot be resolved individually but the overall level of under-determination is reduced. There is also redundancy in the system of equations, since for two isotopic species, the abundance can be estimated in two ways: 111 can be quantified from either NMR data (eq 23a) or MS data (eq 23b), and 011 from different combinations of MS and NMR data (eqs 21a and 21b).

This intuitive example with ^1^H-NMR, ^13^C-NMR and MS data highlights how symbolic calculations can be used to develop a generic framework for integrating isotopic measurements. The calculations are purely based on the fundamental relationships between measurements and the underlying isotopic species. The proposed framework can integrate measurements from any analytical platforms to identify individual species that can be quantified, as well as the combination of species that cannot be resolved individually. This approach thus clarifies the coverage of isotopic space by transforming platform-dependent measurements into (a set of) isotopic species that can be actually quantified. The proposed framework can also consolidate isotopic datasets by integrating quantitative measurements into a single non-linear least squares (NLS) problem. These different aspects are explained in the following sections.

### Mathematical formulation and implementation

Eqs 1–4 can be reorganized into linear isotopomer equations with measurements as parameters. For measurement *a*, this gives:

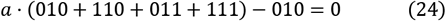

And linear equations can be obtained similarly for all the other measurements. Let *A* be the resulting *m* by *p* matrix, *x* a *p*-length vector of isotopomers, and *b* the right hand side vector of length *m*:

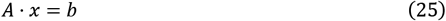

Note that *A* and *b* depend on measurements only. The rows of *A* are linearly dependent as the measurements are normalized to add up to 1.

Eq 25 can be solved by first reducing *A* to its echelon form by Gauss-Jordan elimination:

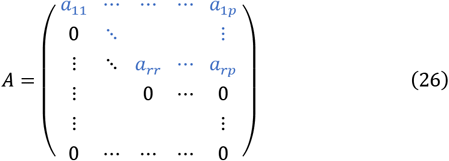

The elements in blue can be non-zero while those in black are all 0. The elements on the main diagonal from *a_11_* to *a_rr_* are strictly non-zero. The number of linearly independent rows defines the *rank* of the matrix, *r*. Two situations can arise depending on the datasets considered:

- If *r < p*, the system is under-determined, with *p-r* free isotopomers. Some of the isotopomers depend only on measurements (we call these *defined* isotopomers), while others also depend on *free* isotopomers. It can also happen that some isotopomers are defined by multiple sets of measurements. This is referred to as measurement redundancy, of which eqs 21a,b and eqs 23a,b are examples;
- If *r* = *p*, the system is just-or over-determined. All isotopomers can be defined uniquely, or in multiple ways in the case of measurement redundancy.

The next step in solving eq 25 is to back-solve the echelon form from *x_r_* to *x_1_*. During echelon reduction and back-solving, the expressions obtained are simplified at each stage using the fact that the measurements from a given method add up to 1. This property was not included in the standard SymPy^21^ module we used to manipulate symbolic expressions so we developed dedicated procedures to solve and simplify these expressions. These can be found in the IsoSolve source code.

Once the vector *x* is obtained as a function of measurements and possibly free isotopomers, measurement redundancy is assessed by substituting isotopomers into the equations defining the measurements, e.g. substituting the solution of eq 25 into eqs 1–4. If in this process, a definition only contains measurements from methods different from the one considered, it is declared redundant.

Formulas for cumomers^22^ (i.e. cumulative isotopomers, which describe sets of isotopomers) and EMUs^17^ involving measurements only are obtained in a similar way. Solutions for isotopomers are substituted into the equations defining cumomers and EMUs and simplified. Defined cumomers and EMUs are then those without free isotopomers in their final formulas.

When the system is under-determined, one point of interest is: which combinations of isotopomers are still measurable, i.e. not dependent on free isotopomers. This question is addressed by exhaustively testing isotopomers that depend on free isotopomers to identify combinations that can be expressed without free isotopomers. During this procedure, combinations involving shorter, already identified combinations are ignored, such that only elementary measurable combinations are identified.

Figure 1 illustrates how this algorithm is implemented in IsoSolve (Figure 1).

**Figure 1.**
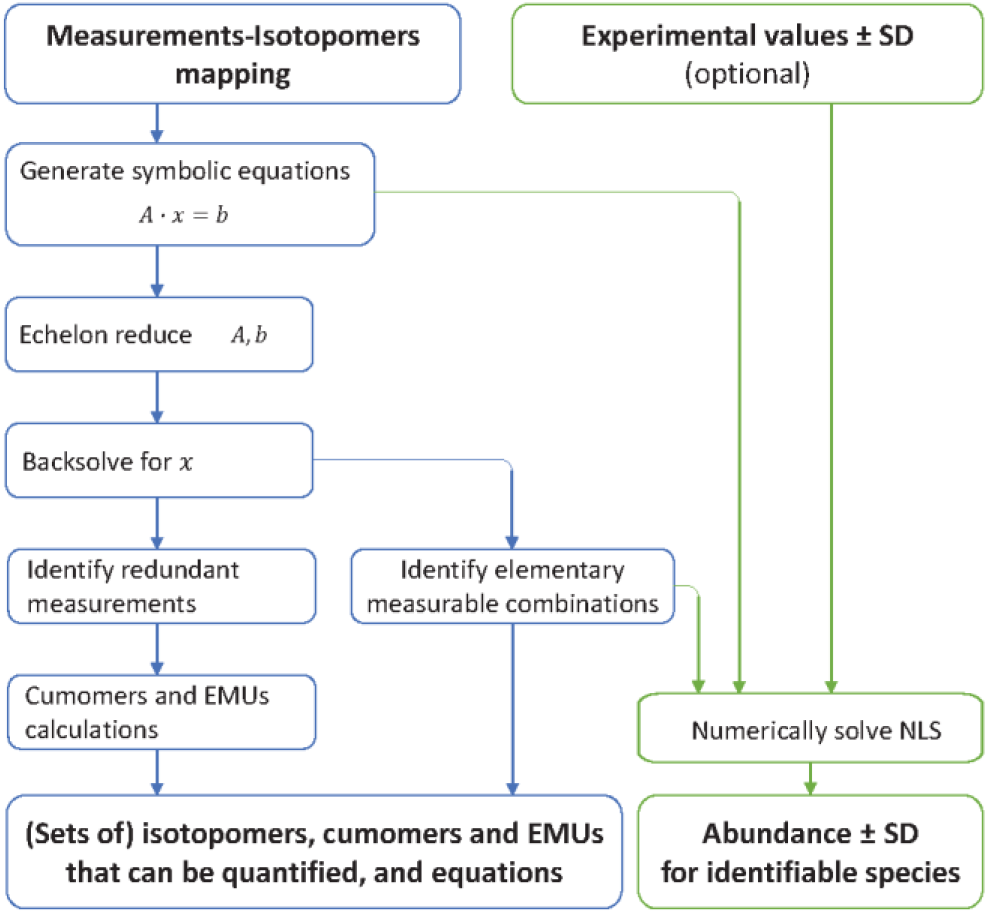
Algorithm implemented in IsoSolve to integrate partial, heterogeneous isotopic measurements. IsoSolve takes as input the relationships between measurements and isotopomers, as defined in Figure 2, to identify the (sets of) isotopomers, cumomers and EMUs that can be quantified individually and produce the corresponding equations (steps framed in blue). When numerical values of measurements are also provided, IsoSolve determines the abundance of all identifiable species (optional steps in green).

A useful feature of IsoSolve that is not directly related to symbolic equation resolution is that experimental data can be consolidated by solving the appropriate NLS problem (Figure 1). The equations in this problem are identical to those defining measurements, such as eqs 1–4. The numerical solution minimizes the cost function *T(x)* defined as the sum of squared residuals. Each residual *u*_*i*_(*x*) is calculated as the difference between experimental measurement *i* 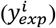 and its value calculated from estimated isotopomers 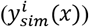, normalized by their respective SDs 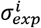, provided by the user):

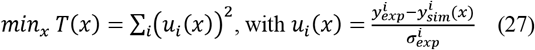

Obvious constraints of necessarily non-negative values that add up to 1 are applied to the solution. This NLS problem with inequality and equality constraint is solved using the NLSIC algorithm^23^. A chi-square test is performed to determine if the fit is satisfactory (based on a 95% confidence threshold). Discrepancies indicate inconsistencies between the different datasets. Finally, IsoSolve estimates the precision of the abundance of each isotopomer, cumomer and EMU by propagating measurement uncertainties. Considering a linearized relationship between small variations in the residual vector Δ*u* and induced variations in the solution vector Δ*x*:

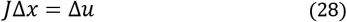

Where *J* is the Jacobian matrix of partial derivatives *∂u*/*∂x*, the covariance matrix cov(*x*) is related to a given covariance matrix cov(*u*) as:

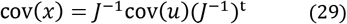

Given that *J* is not necessarily invertible, we use a singular value decomposition (SVD) of *J* = *UD*(*s*)*V*^*t*^ where *U* and *V* are orthogonal matrixes and *D(s)* is a diagonal matrix with a vector *s* of strictly positive elements defining the main diagonal. The length of this vector is equal to the rank of *J*, *r_J_*. Since the residuals are scaled by SDs, their covariance matrix is expected to be an identity matrix and the final expression simplifies to:

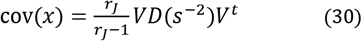

Similar to Bessel’s correction, the factor *r*_*J*_/(*r*_*J*_ − 1) ensures that the estimator is not biased. The SDs of *x* are simply the square roots of the elements on the main diagonal of cov(*x*). For sake of brevity, the fact that *x* is constrained to sum to 1 has been omitted from the above description; this constraint is however taken into account in IsoSolve.

### Clarifying the isotopic coverage of alanine for individual and combined methods

Combining different analytical methods should improve the coverage of isotopic space. As a first step, we used this workflow to clarify the isotopic information provided by combining a broad range of (NMR and/or MS) methods. Based on the literature^10–11^, ^13^ ^19–20^, ^24–25^, we defined a list of eight datasets (D1-D8) that can be obtained for alanine (Figure 2). Even though it only contains three carbon atoms, no single method can completely cover alanine's isotopic space by itself.

**Figure 2.**
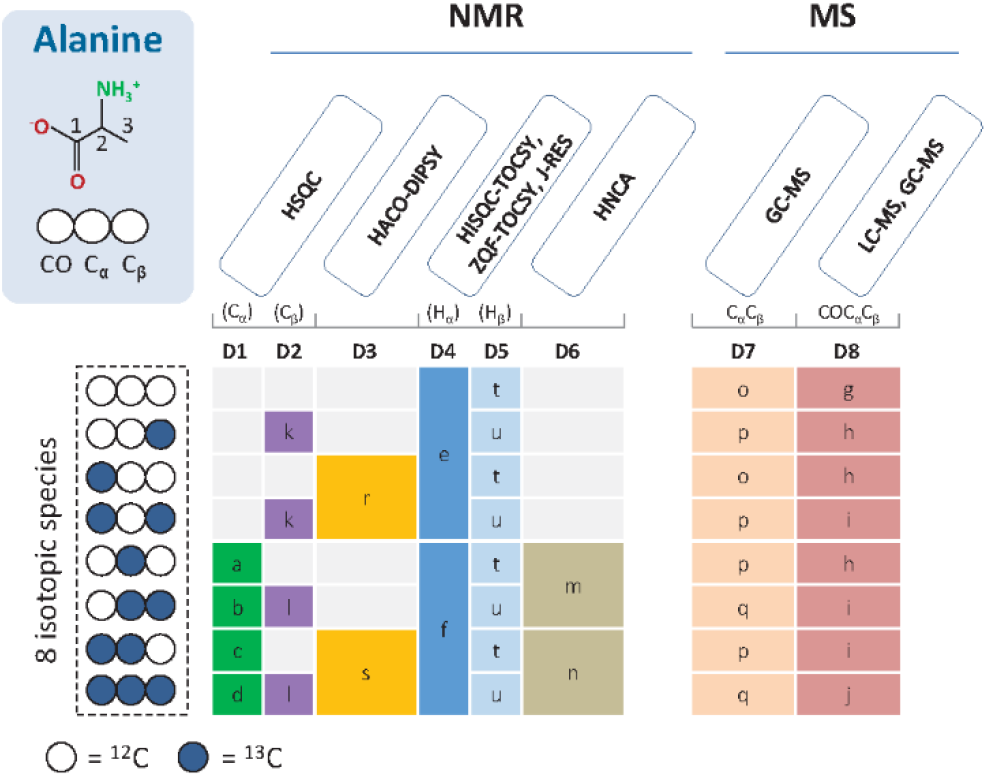
Isotopic measurements on alanine by eight NMR and MS methods. The eight isotopic species of alanine are shown on the left, with white and blue circles representing ^12^C and ^13^C atoms, respectively. Methods providing the same information are grouped together (e.g. ZQF-TOCSY and J-RES NMR experiments). For each dataset(D1-D8), each group of measurements is shown by a specific color, and the letters refer to the (sets of) species that are quantified relative to the amount of all species present in the corresponding group.

We evaluated all 255 possible combinations of datasets. The combinations were evaluated based on the following metrics: number of individually quantifiable isotopomers and cumomers, number of EMUs for which all isotopologues can be quantified, number of redundant measurements, and information gainedfrom the proposed integrative framework (i.e. number of additional quantifiable isotopomers).

The results are summarized in Figure 3. In most situations, integrating different datasets improves the coverage of isotopic space, though some combinations do not provide any new information (e.g. D2+D3+D4+D5, combination #127). Importantly, 34% of the combinations (87/255) provide complete coverage of the isotopic space of alanine (i.e. quantify all its isotopomers, cumomers and EMUs), with different degrees of redundancy (from 0 to 10 redundant measurements). This is only possible if the combined dataset includes both NMR and MS data, highlighting the complementarity of the two techniques. This analysis shows that all the isotopomers of alanine can be quantified with as few as three datasets.

**Figure 3.**
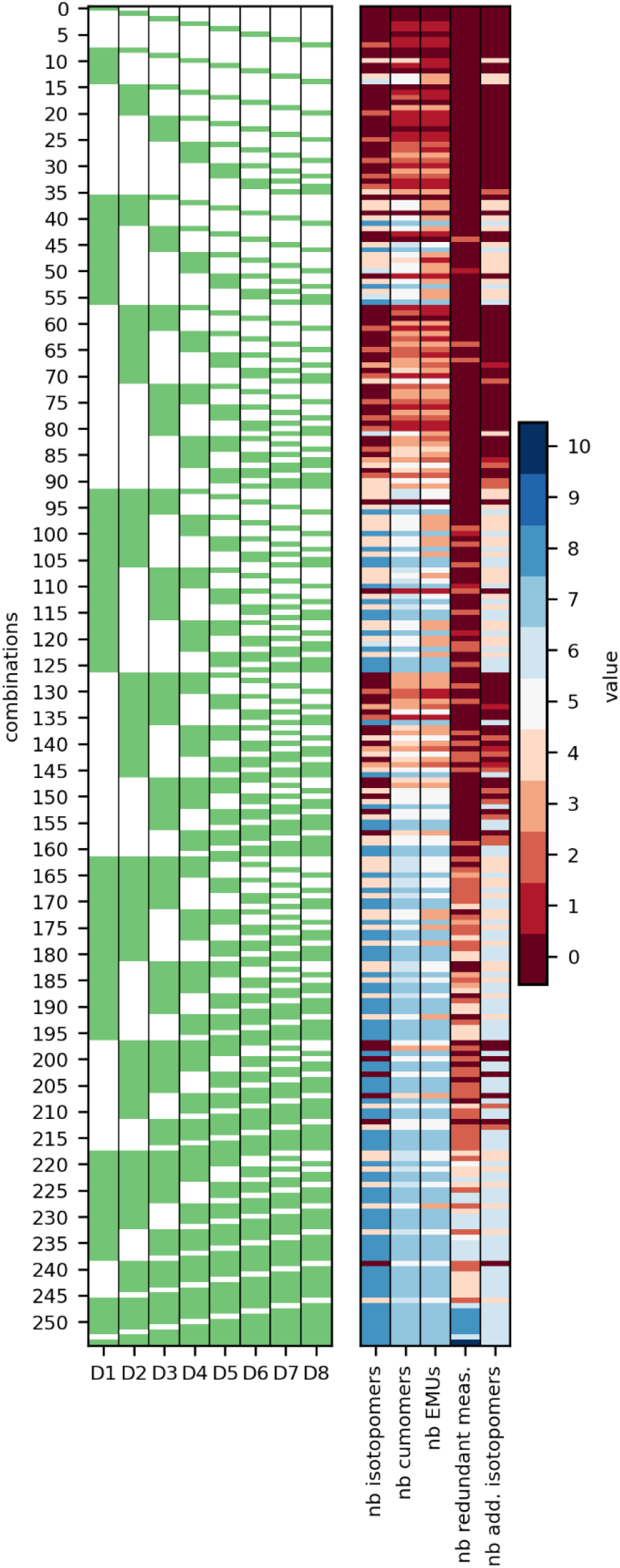
Isotopic information for alanine obtained by integrating different datasets. For each of the 255 possible combinations of datasets (left panel, where each line represents a combination, with included datasets shown in green), the following metrics were calculated (right panel): number of individually quantifiable isotopomers, cumomers and EMUs, number of redundant measurements, and number of additional isotopomers quantifiable only thanks to data integration.

Indeed, combining LC-MS and NMR HSQC data (with C_α_ and C_β_ signals, D1+D2+D8, combination #41) leads to the following solution (where the letters refer to the measurements shown in Figure 2):

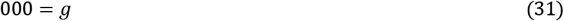

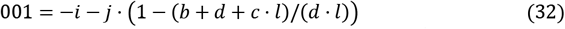

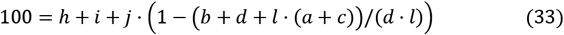

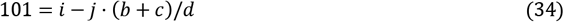

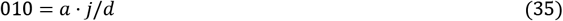

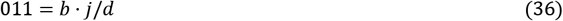

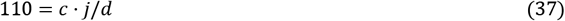

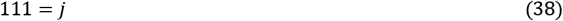

As well as improving isotopic coverage, this analysis may thus be used to guide experimental design by identifying the best combination of analytical methods and datasets to quantify a given set of isotopic species. As demonstrated here, the complementarity of any technique is readily evaluated, hence providing guidance for future analytical developments.

### Isotopic coverage of proteinogenic amino acids

Following the same approach, we used IsoSolve to clarify the current isotopic coverage of proteinogenic amino acids by determining the number of individually quantifiable isotopomers, cumomers and EMUs. Four amino acids are lost during protein hydrolysis (cysteine, tryptophan, glutamine, and asparagine) and cannot be detected. Data integration was thus carried out for the remaining 16 proteinogenic amino acids.

Figure 4 highlights the heterogeneity of isotopic coverage for the different amino acids. While complete isotopic coverage is achievable for amino acids containing up to three carbon atoms (glycine, serine and alanine), the coverage progressively decreases as the number of carbon atoms increases. Isotopomer coverage was 12-31% for C_4_-amino acids (aspartate, threonine), 6-12% for C_5_-amino acids (glutamate, methionine, proline, valine), 3-6% for C_6_-amino acids (arginine, histidine, leucine, isoleucine, lysine), and about 1% for C_9_-amino acids (phenylalanine and tyrosine). As expected, the coverages are higher for cumomers (100% for C_2_ and C_3_, 40-53% for C_4_, 23-42% for C_5_, 6-24% for C_6_, 2-3% for C_9_) and for EMUs (100% for C_2_ and C_3_, 27-40% for C_4_, 16-32% for C_5_, 5-21% for C_6_, 1-2% for C_9_) than for isotopomers. Overall, this framework provides a clear audit of the isotopic information that can actually be measured on proteinogenic amino acids.

**Figure 4.**
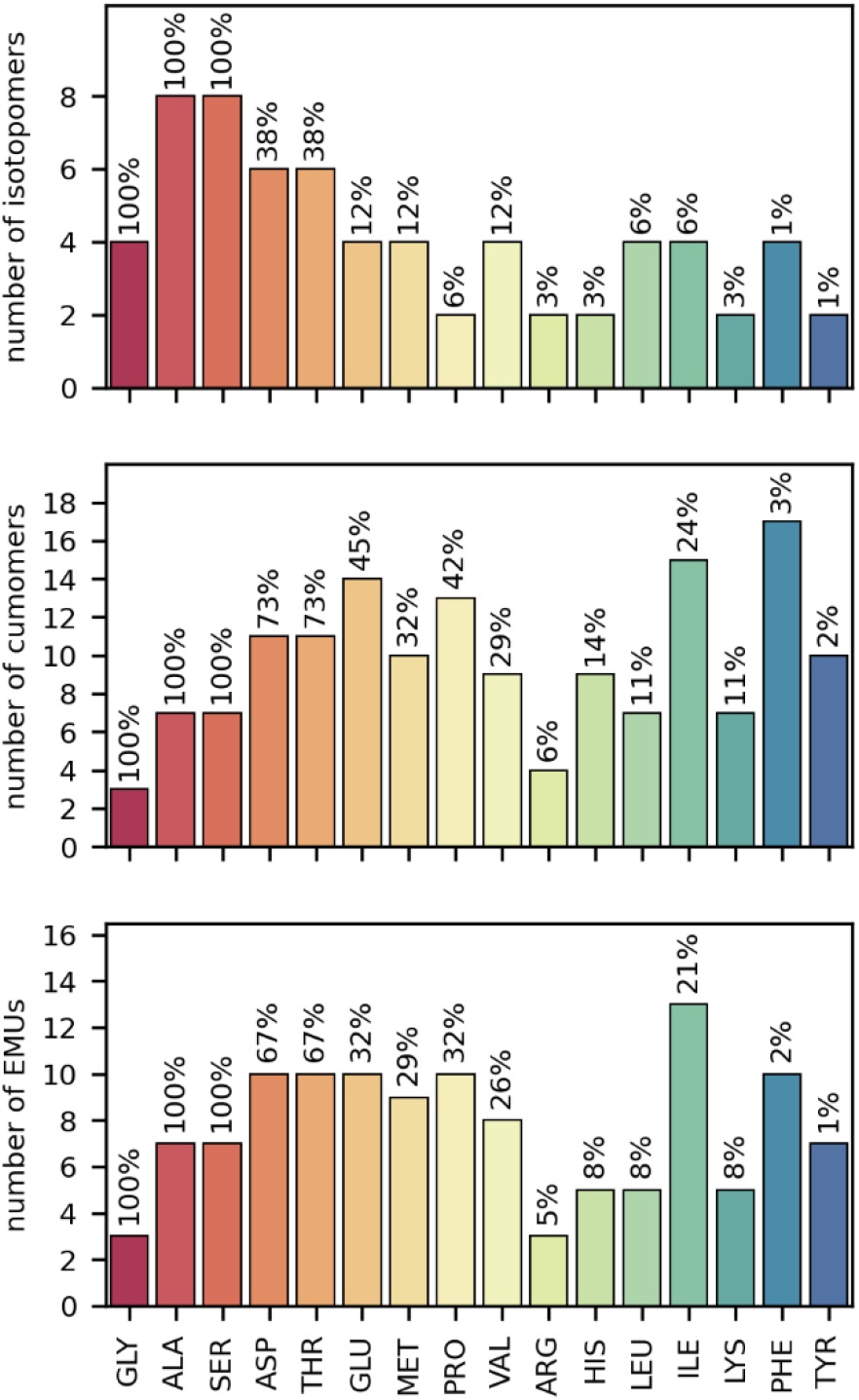
Isotopic coverage of proteinogenic amino acids. Number of isotopomers, cumomers and EMUs that can be quantified individually for each amino acid by integrating available datasets. The respective proportions of isotopic forms that can be quantified are shown above the bars.

### Data integration and consolidation

To quantitatively evaluate the proposed integrative framework, we produced a reference sample of ^13^C-labeled proteinogenic amino acids with controlled and predictable labeling patterns in which all isotopic species are present in equal amounts^9^.

This sample was analyzed by NMR (ZQF-TOCSY, HSQC, HNCA and HACO-DIPSY experiments) and MS (GC-MS and LC-MS), yielding eight independent datasets (D1-D8 in Figure 2) containing a total of 21 isotopic measurements for alanine (Supporting information S1). For all combinations of datasets identified as providing complete isotopic coverage of alanine (Figure 3), we used IsoSolve to determine the abundance of each isotopomer, cumomer and EMU. The results obtained for each combination were evaluated using two quantitative metrics^19^: accuracy (defined as the mean error and calculated from the differences between theoretical and measured abundances) and precision (defined as the mean standard deviation of the measured abundances).

All the isotopomers, cumomers and EMUs of alanine were indeed quantified for all the combinations considered. The accuracy and precision of the results depended on the datasets included (Figure 5).

**Figure 5.**
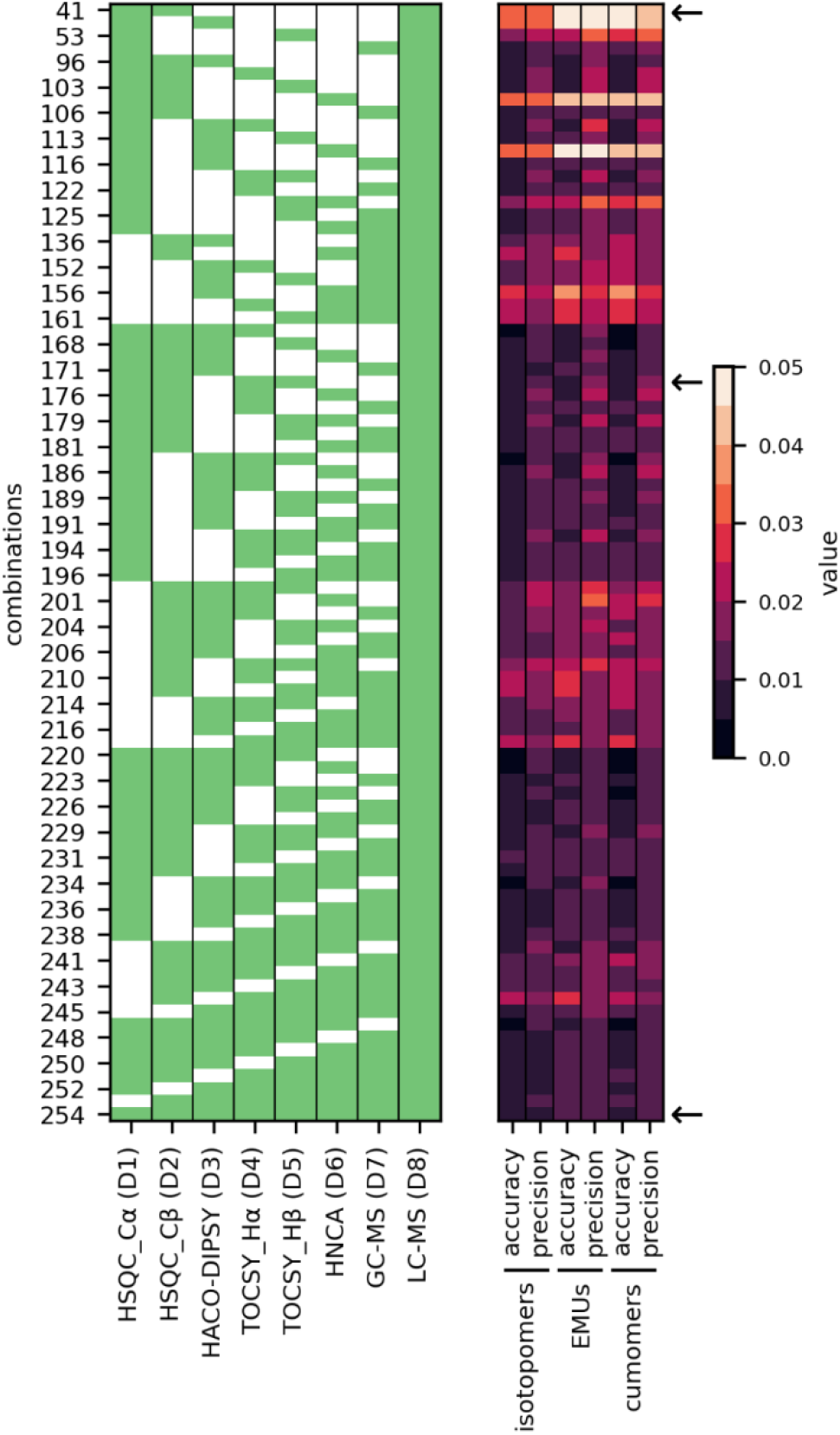
Summary of data integration results for all dataset combinations that completely cover the isotopic space of alanine. The accuracy and precision of isotopomer, cumomer and EMU quantifications (right panel) were determined for each combination of datasets (left panel). Combinations discussed in the text and detailed in Figure 6 are indicated by an arrow.

Regarding isotopomers for example, integrating HSQC and LCMS datasets (D1+D2+D8, combination #41) led to an accuracy and precision of 0.033. Adding the TOCSY NMR dataset (with H_α_ and H_β_ signals, D1+D2+D4+D5+D8, combination #174) improved both metrics (accuracy = 0.006, precision = 0.014) and results were further improved when all datasets were combined (D1-8, combination #254, accuracy = 0.006, precision = 0.009). Similar trends were observed for cumomers and EMUs (Figure 5).

A detailed analysis of the integration results reveals that precision and accuracy also depend on the isotopic species considered (Figure 6). Combining HSQC and LC-MS data is sufficient to reliably quantify 6 of the 8 isotopomers of alanine. Isotopomers 000 (accuracy = 0.002, precision = 0.009) and 011 (accuracy = 0.020, precision = 0.020) are for instance well resolved, but isotopomers 001 (accuracy = −0.060, precision = 0.070) and 100 (accuracy = 0.102, precision = 0.082) remain poorly resolved. Adding the TOCSY dataset significantly improves quantification for the two latter species (001: accuracy = 0.018, precision = 0.022; 100: accuracy = −0.008, precision = 0.025). The most reliable results were obtained by integrating all the datasets (001: accuracy = 0.018, precision = 0.013; 100: accuracy = −0.007, precision = 0.010). Here again, similar trends were observed for cumomers and EMUs (Figure 6).

**Figure 6.**
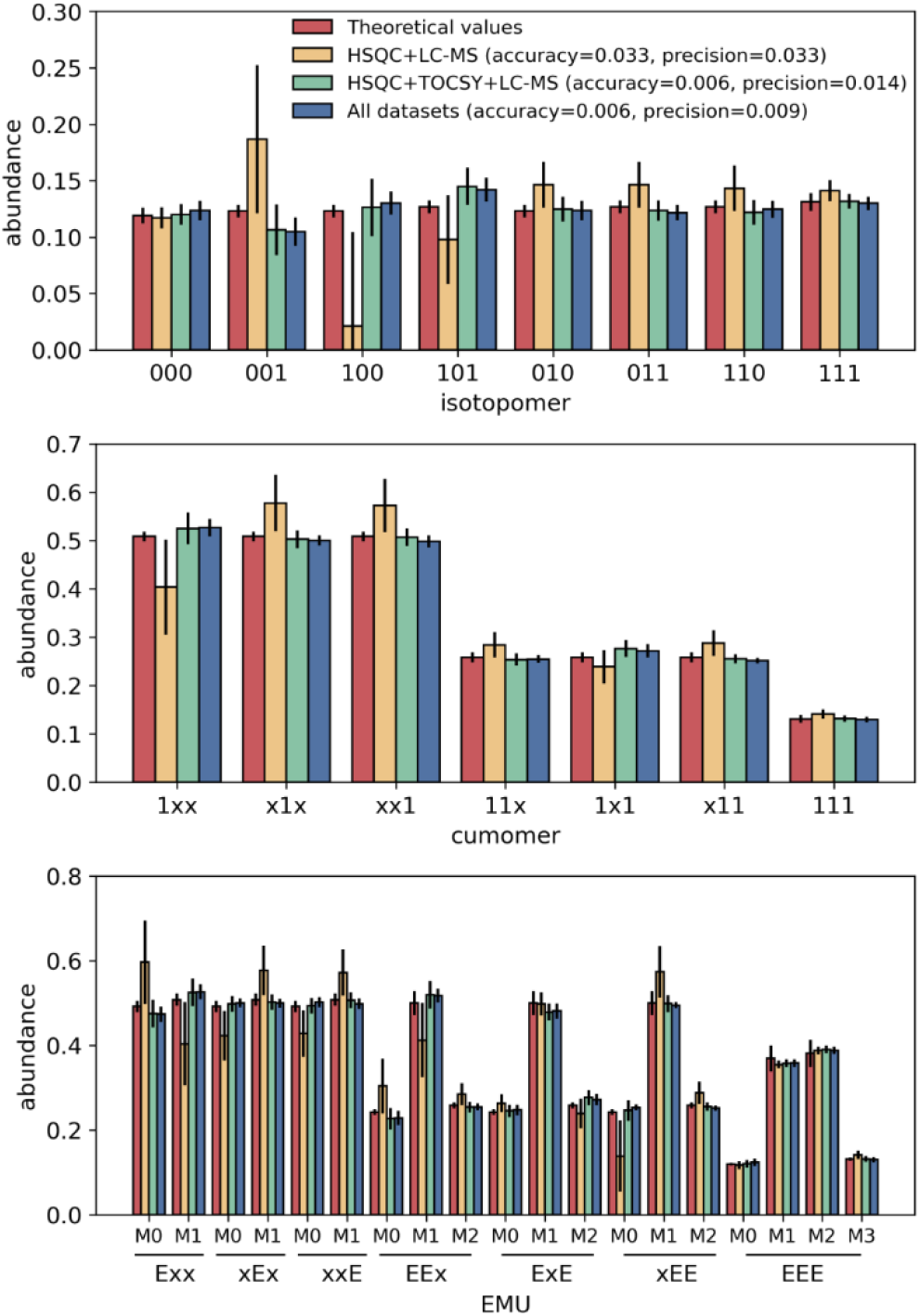
Detailed quantification results for three different combinations of datasets. The abundance of each isotopomer (upper panel), cumomer (middle panel) and EMU (lower panel) was estimated by integrating different datasets (HSQC+LC-MS, orange bars; HSQC+TOCSY+LC-MS, green bars; all datasets, blue bars), and experimental values were compared to the theoretical abundances in the reference sample (red bars). ^12^Cand ^13^C-atoms are represented by 0s and 1s, respectively; *x* stands for “0 or 1”, and *E*s denote the atoms contained in the corresponding EMU. Error bars correspond to ± one standard deviation.

In combinations with measurement redundancy, the consistency of the different datasets can be evaluated using a chi-square test. For instance, when all datasets were combined (D1-8, combination #254, 10 redundant measurements), the chi-square test confirmed that all the measurements were consistent (p-value = 0.994). When some of these measurements were artificially altered, e.g. *b* changed from 0.2533 to 0.0533 and *c* from 0.2475 to 0.4475, the p-value decreased to 2×10^−16^, highlighting the inconsistencies between the redundant measurements. This illustrates how the proposed approach can be used to identify biased measurements to be checked before interpretation.

These results confirm that integrating additional datasets improves both the accuracy and the precision of isotopomer quantification, very likely because of the high degree of redundancy (up to 10 redundant measurements) which reduces the impact of experimental noise and the potential biases of individual measurements. Overall, all isotopomers could be reliably quantified using a wide variety of data combinations, with a high accuracy and precision in most situations.

## CONCLUSION

The complementarity of different (MS and/or NMR) approaches dedicated to isotopic analyses is often highlighted, but the lack of a generic integrative framework able to deal with heterogeneous, partial isotopic measurements has meant that this has never been evaluated in detail. The proposed framework fills this conceptual gap by allowing any type of isotopic measurements (MS, MS/MS, ^1^HNMR, ^13^C-NMR, ^15^N-NMR, etc) to be included. The framework is agnostic to the analytical platform, the labeled element, the tracer isotope, and the molecule. It can also be applied to double-labeling approaches (e.g. ^13^C and ^15^N). This framework has been implemented as an open source Python program, IsoSolve, which is available as a command-line interface and as a Python library to streamline its integration into existing data analysis pipelines.

Using amino acids as an example application, we have demonstrated that this framework can i) clarify the actual coverage of isotopic space by identifying the (sets of) species that can actually be quantified, ii) improve this coverage by increasing the number of isotopic species that can be quantified individually, iii) evaluate the complementarity and redundancy of different techniques, iv) consolidate isotopic datasets by evaluating their consistency, identifying biased measurements, and reducing the impact of measurement noise, v) support experimental design by identifying the most relevant methods to quantify a given set of isotopic species, and vi) guide future analytical developments.

Our framework connects measurements to the chemical structure of compounds and to their formal representation in isotopic models of metabolism, hence assisting both model-free and model-based data interpretation. This framework may thus support structural investigations (e.g. metabolite identification, spectra annotation, validation of MS/MS fragmentation patterns, development of standardized databases for deposition of isotopic datasets based on isotopically-resolved InChIs) as well as functional investigations of metabolic systems (e.g. experimental design, data consolidation, conversion between isotopic representations in ^13^C-fluxomics workflows). It should also make isotope labeling experiments more accessible to the wider biological community.

Beyond isotopic studies, this framework may prove equally valuable in other fields dealing with the analysis of combinatorial states of biological entities. This is the case in proteomics for instance, for the analysis of post-translational modifications (e.g. phosphorylation or acetylation), with the ultimate objective of determining the complete distribution of each of the 2^n^ forms of a protein with n-modification sites. Partial information on these distributions can be obtained by MS and NMR^26^, and these datasets can be integrated using IsoSolve following the principles described in this article.

## AUTHOR INFORMATION

### Notes

The authors declare no competing financial interest.

## ACKNOWLEDGMENT

The authors thank MetaboHub-MetaToul (Metabolomics & Fluxomics facilities, Toulouse, France, http://www.metatoul.fr), which is part of the French National Infrastructure for Metabolomics and Fluxomics (www.metabohub.fr), funded by the ANR (Metab-oHUB-ANR-11-INBS-0010), for access to NMR and MS facilities. JCP is grateful for funding from INSERM for his temporary full-time researcher position.

